# Evolutionary history limits species’ ability to match color sensitivity to available habitat light

**DOI:** 10.1101/2021.10.29.466507

**Authors:** Matthew J. Murphy, Erica L. Westerman

## Abstract

The spectrum of light that an animal sees – from ultraviolet to far red light – is governed by the number and wavelength sensitivity of a family of retinal proteins called opsins. It has been hypothesized that the spectrum of light available in an environment influences the range of colors that a species has evolved to see. However, invertebrates and vertebrates use phylogenetically distinct opsins in their retinae, and it remains unclear whether these distinct opsins influence what animals see, or how they adapt to their light environments. Systematically utilizing published visual sensitivity data from across animal phyla, we found that terrestrial animals are more sensitive to shorter and longer wavelengths of light than aquatic animals, and that invertebrates are more sensitive to shorter wavelengths of light than vertebrates. Controlling for phylogeny removes the effects of habitat and lineage on visual sensitivity. Closed and open habitat terrestrial species have similar spectral sensitivities when comparing across the Metazoa, and deep water animals are more sensitive to shorter wavelengths of light than shallow water animals. Our results suggest that animals do adapt to their light environment, however the invertebrate-vertebrate evolutionary divergence has limited the degree to which animals can perform visual tuning.

## 1. Introduction

Animals use vision for many tasks, including finding prey, avoiding toxic animals and plants, identifying predators, assessing mate quality, and navigating their environment [1–5]. In many cases, the objects of interest to the animal need to be distinguished from the background [6,7]. For example, food that does not contrast with the background is harder for foragers to detect than food that does contrast with the background [8–12]. Signals that contrast with background colours and patterns are also used for mating displays [13–18]. Furthermore, many species’ body colour patterns have evolved to be simultaneously cryptic to predators while conspicuous to intended receivers [19–21]. Finally, contrasting colours can improve animals’ ability to learn the meaning of signals, as when chicks learn more quickly to avoid bitter, aposematically coloured food [2,3,22].

An animal’s ability to detect visual information depends upon the colour and amount of light in its habitat, otherwise known as the *light environment* [6,23,24]. For example, red and blue light are filtered out by chloroplasts, lending forests and estuarine environments a yellow-green cast [24–26]. Likewise, the water column progressively filters red and UV light [23,24]. Animals’ signalling behaviours, choice of microhabitat, and visual physiology are thus expected to co-evolve to suit their light environment [6].

Sighted species’ photoreceptors (the light-absorbing neurons which enable vision) are theorized to have undergone adaptation to best absorb the light most often present in their environments [23]. This process, called *visual tuning*, is made possible by both filtering pigments [27–35] and differences in the amino acid sequence or 3-dimensional shape of photosensitive proteins called *opsins* [23]. Visual tuning has been found to shift wavelengths of maximum sensitivity in species as diverse as birds, fish, and mammals [23,36,45,46,37–44]. Although the effect of light environment on vision has been extensively studied in fish [47,48], a systematic study of visual tuning in terrestrial animals has not yet occurred; neither have terrestrial animals been systematically compared to aquatic species. Both aquatic and terrestrial animals are found in a variety of light environments, and multiple phyla have independently made the water-to-land habitat transition. Additionally, studies of animals which transition from aquatic larvae to terrestrial adults have found that these species change their visual pigment expression patterns in a manner that matches their changing light environment [49–53]. Understanding whether phylogeny constrains the extent of visual tuning, particularly during these water-to-land transitions, is critical for understanding the evolutionary ecology of animal vision.

If opsin tuning faces phylogenetic constraints, the evolutionary history of animal vision may have shaped the degree to which different phyla have adapted to their light environments. The types of opsins differ between chordates and other phyla [54]. Chordates use c-opsins in cilia-bearing photoreceptors to transduce photons into vision, while non-chordate animals use r-opsins in rhabdomere-bearing photoreceptors; no animals have been identified that use both c- and r-type photoreceptors for vision [55]. Phylogenetic analyses reveal that c- and r-opsins diverged 400 million years ago and were likely both present in the urbilaterian, with r-opsins closely related to the melanopsins used by chordates for non-visual tasks [55]. The r-opsin/ c-opsin divergence may have given rise to different degrees of tuning between chordates and non-chordates.

The diverse habitats in which animals live, combined with the long evolutionary history of visual pigments, leads to several questions. First, have transitions from aquatic to terrestrial habitats influenced the spectra of light that animals can see? And, are differences in the spectra that animals can see associated with the c-opsin/ r-opsin divergence? Second, do animals that live in visual environments that filter red and blue light, such as closed-canopy forests and estuarine habitats, see colours more similarly to each other than to open terrestrial or freshwater aquatic environments, in which colours are less (if at all) strongly filtered? And, if there is an effect of habitat greenness, is this effect outweighed by phylogeny?

To answer these questions, we performed a phylogenetically weighted systematic analysis of the maximum and minimum wavelength of visual sensitivity, as well as the range of visual sensitivity, across animals.

## (2) Materials and method

### Paper selection

We conducted Google Scholar searches in October 2017 and January 2018. Our first search used the search pattern *“visual pigment” OR opsin OR “absorbance spectrum” “λ max” -human -man -men -woman -women - ”Homo sapiens” -disease -regeneration*. We conducted a second Google Scholar search using the search pattern *visual pigment, opsin sensitivity, absorbance spectrum*. For both searches, we excluded citations and patents.

We reviewed candidate articles using a three-step process. First, we screened by title and abstract to identify original research articles and review papers that examined animal visual physiology. We then screened articles to determine if they used microspectrophotometry, electrophysiology, pigment extraction, or *in vitro* mRNA expression followed by spectrophotometry, and that they measured visual sensitivity or visual pigment absorption from at least two animals. Finally, we only kept articles which used animals that were wild-caught or reared in full-spectrum light conditions, to avoid any effects of artificial lighting on visual sensitivity [56,57].

For review articles, we determined whether the authors had included measurements of the mean wavelength of peak sensitivity (λ_max_) of some population in the article’s figures or tables. We downloaded the corresponding primary sources and filtered them using the process described above.

### Visual pigment sensitivity data

We recorded the following data for each species of each paper that passed our filters: 1) mean wavelength of peak sensitivity (λ_max_) for each visual pigment measured; 2) number of animals measured (*n*); 3) standard deviation of the mean λ_max_ (SD) (when available); and 4) where animals were caught (when available). We calculated sampling error for visual pigments when possible.

### Habitat data

We used standardized data sources to classify each species by habitat. Sources included field guides [58–60], public databases (BugGuide, <bugguide.net>, Butterflies and Moths of North America, <butterfliesandmoths.org>, FishBase <fishbase.org>, SealifeBase <sealifebase.org>, IUCN Redlist <iucnredlist.org>) and online encyclopaedias including Animal Diversity Web (<animaldiversity.org>) and Encyclopedia of Life (<eol.org>). After first classifying species as terrestrial or aquatic, we then defined terrestrial sub-habitats: rainforest, forest, woodland, shrubland, grassland, and desert. We recategorized these habitats into three habitat types based on canopy density. Rainforest and temperate forest were designated as “closed” habitats. Woodland was considered to have “intermediate” canopy density [25]. Shrubland, grassland, and desert were classified as “open” habitats.

Aquatic habitats included river, stream, pond, lake, coastal, estuarine, open-water marine, bottom-dwelling marine, abyssopelagic, abyssodemersal, bathypelagic, and bathydemersal habitats. We recategorized these habitats into two habitat types based on salinity. River, stream, pond, and lake habitats were considered “freshwater” habitats; while coastal, estuarine, open-water marine, and bottom-living marine habitats were “marine” habitats. Animals considered “coastal” were those described as living in water along the coast, near shore, or in estuaries. We also recategorized these habitats into two habitat types based on whether light was abundant or not. Abyssopelagic, abyssodemersal, bathypelagic, and bathydemersal habitats receive little or no sunlight due to their depth in the water column and were considered “lightless” habitats. Species that were considered by our sources as deep-water species were also considered species that lived in “lightless” habitats. All other habitats were considered “lit” habitats. Finally, we used FishBase, SealifeBase or field guides to identify the minimum and maximum depths for each species. We then used these data to calculate average depth per species (D_average_ = (D_max_+D_min_) *2^-1^).

### Phylogenetic control

To control for the effect of evolutionary relatedness on visual sensitivity we built a phylogenetic tree of all animals in our analysis (see the electronic supplementary material: figure S1). We used the function tnrs_match_names in the R package *rotl* [61] to acquire data from the Open Tree of Life database (<tree.opentreeoflife.org>) for each of the species represented in our regression, and to generate a phylogenetic tree using default arguments and excluded species flagged as *incertae cedis;* i.e., with uncertain phylogenetic position (25 species) and species which had no sequencing data in the Open Tree of Life database (6 species). We created an induced subtree with the resulting data using the function tol_induced_subtree in the package *rotl*.

Trees produced using *rotl* are unrooted, without branch lengths, and sometimes with unresolved polytomies. We used the R packages *phytools* [62] and *ape* [63] to resolve these issues. We used the root function in *ape* to root the tree using *Saccharomyces cerevisiae* (ottid: 5262624) from <tree.opentreeoflife.org>) as the outgroup. We computed branch lengths using the compute.brlen function in *ape* with default arguments. Finally, we randomly resolved polytomies using the multi2di function in *ape* with default parameters. Subtrees of the primary tree were constructed as needed using the drop.tips function in *ape*.

### Statistical Analyses

To determine whether longest λ_max_, shortest λ_max_, and range of λ_max_ followed the normal distribution, we used the shapiro.test function in R. To determine whether the variances of longest λ_max_, shortest λ_max_, and range of λ_max_ differed between broad habitat type (aquatic or terrestrial) or lineage (invertebrates or vertebrates) we used levene.test function in R.

To determine whether there was an effect of broad habitat type or lineage on the longest λ_max_, shortest λ_max_, and range of λ_max_, we used the glm function in R to construct generalized linear models with the formula λ_max_ ~ broad habitat * lineage.

To determine whether phylogeny could explain extant differences in longest λ_max_, shortest λ_max_, and range of λ_max_ between broad habitat type or lineage, we constructed phylogenetically controlled linear models using the phylolm function in the *phylolm* [64] package for R with the formula λ_max_ ~ broad habitat * lineage with a bootstrap of 100. Since we had to exclude 31 species from our phylogenetic tree, we first ran the glm described above with the trimmed data set, and then compared those results to the results of our phylolm models. For these models, we used the overall phylogenetic tree and our trimmed dataset.

We then subset our overall dataset for terrestrial animals and aquatic animals, re-tested for normal distributions and variances, and conducted a set of statistical analyses specific to terrestrial or aquatic animals. To determine whether there was an effect of terrestrial habitat type (closed, intermediate, or open) or lineage on longest λ_max_, shortest λ_max_, and range of λ_max_, we used the glm function with the formula λ_max_ ~ terrestrial habitat type * lineage. However, since there were only 2 species that were intermediate habitat specialists, and they had similar visual spectra to those in closed canopies, we combined closed and intermediate habitat treatments into a single treatment, closed_intermediate, and re-ran the generalized linear models described above using the new habitat treatment levels (closed_intermediate vs open).

We examined the effects of depth and habitat on visual sensitivities of aquatic animals. To determine whether there were effects of minimum, maximum, or average depth of habitat on longest λ_max_, shortest λ_max_, and range of λ_max_ among aquatic species, we constructed linear models using the lm command with the formula λ_max_ ~ depth. To determine whether phylogeny could explain extant differences in minimum, maximum, or average depth of habitat among longest λ_max_, shortest λ_max_, and range of λ_max_ among aquatic species, we first re-ran the linear models without the species excluded from our phylogenetic tree, and then constructed phylogenetically controlled linear models using the phylolm function in the *phylolm* package for R with the formula λ_max_ ~ depth with a bootstrap of 100. For these models, we used a subtree of our overall phylogenetic tree (see above), which omitted all terrestrial species.

Finally, we subset our overall dataset for open terrestrial animals and non-deep water aquatic animals (both freshwater and coastal) and conducted the following analyses. To assess whether the visual systems of animals in open terrestrial habitats were more similar to the visual systems of animals in open water habitats (coastal or freshwater) than those of closed terrestrial habitats we compared the longest λ_max_, shortest λ_max_, and range of λ_max_, of species in coastal-aquatic, freshwater-aquatic, terrestrial-closed, and terrestrial-open habitats using the kruskal.wallis function in R. Following this, we performed a pairwise (Steel-Dwass) test using the dscfAllPairs function in the R package *PCMCRplus* [65]. To determine whether phylogeny could explain extant differences in longest λ_max_, shortest λ_max_, and range of λ_max_ between animals living in these four habitat types we re-ran the above analyses only using species in our phylogenetic tree, and then constructed phylogenetically controlled linear models using the phylolm function in the *phylolm* package for R with using the following formula with a bootstrap of 100: λ_max_ ~ coastal-aquatic + freshwater-aquatic + terrestrial-closed and λ_max_ ~ coastal-aquatic + freshwater-aquatic + terrestrial-open.

## (3) Results

Our dataset included 1,114 opsins from 446 species, extracted from a total of 156 articles (See the electronic supplementary materials: table S1). Of these, 868 opsins were recorded from 355 aquatic species, and 246 opsins were recorded from 91 terrestrial species. Our data were not normally distributed (Shapiro-Wilk test: longest λ_max_: p < 0.05, W = 0.94; shortest λ_max_: p < 0.05, W = 0.83; range of λ_max_: p < 0.05, W = 0.70). Shortest, but neither longest nor range of λ_max_ were found to have equal variances when compared across broad habitat and lineage (Levene’s test: longest λ_max_: p = 0.16, t = 1.74; shortest λ_max_: p < 0.05, t = 25.28; range of λ_max_: p < 0.05, t = 21.62).

### (a) Terrestrial species were maximally sensitive to longer wavelengths of light than aquatic species

Terrestrial species were maximally sensitive to longer wavelengths of light than aquatic species, independent of opsin type (GLM, n=433: habitat p = 3.83*10^-8^, t = 5.600, lineage p = 0.309, t = −1.019; interaction: p = 0.595, t = −0.532; λ_max_ longest long-wavelength terrestrial species: 535±41.6 nm, aquatic species: 506±30.6 nm, invertebrate species: 513±38.9 nm, vertebrate species: 512±33.0 nm) (figure 1a).

**Figure 1.**
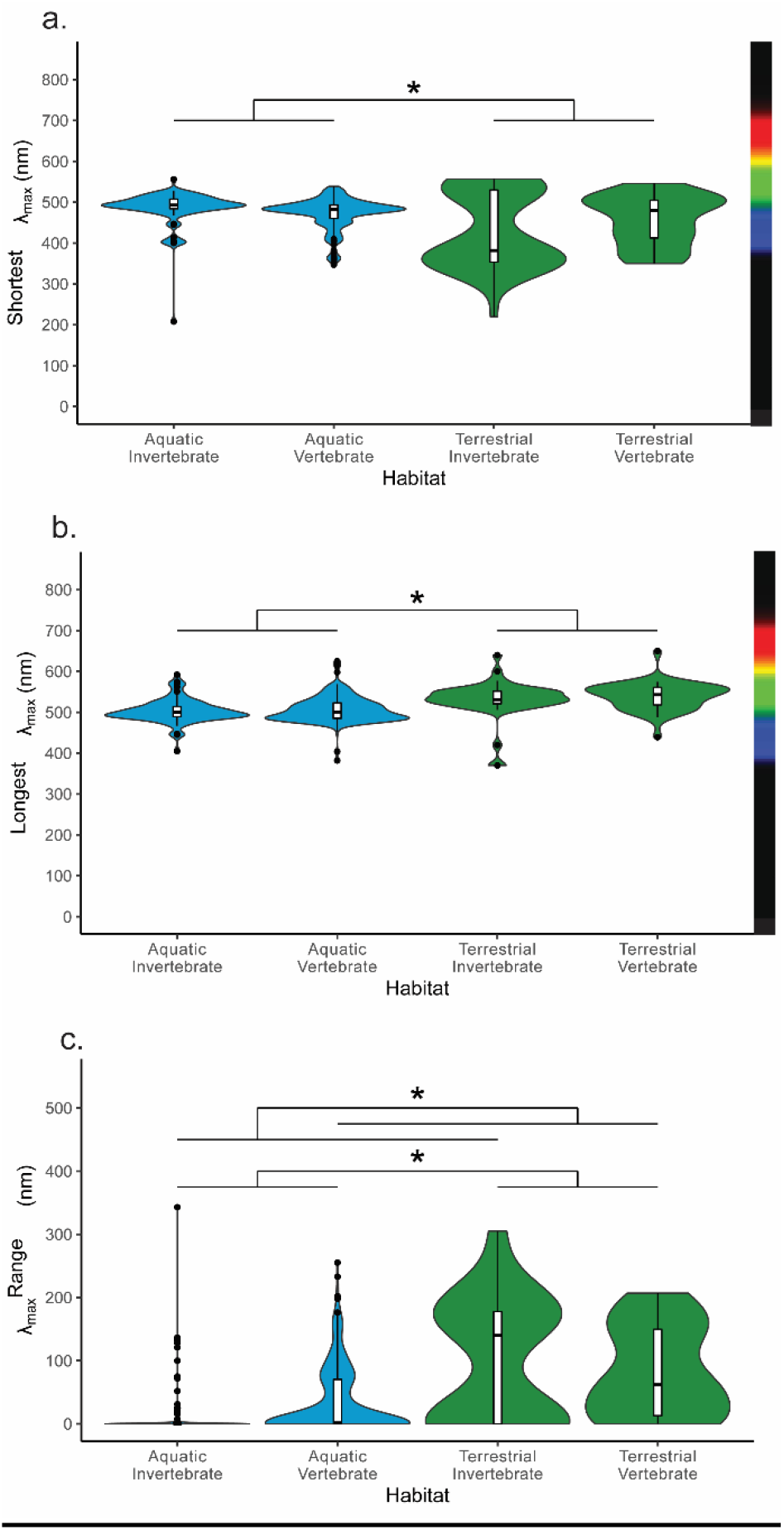
Effect of coarse habitat and lineage on mean visual pigment sensitivity, prior to phylogenetic control. A. Shortest opsin: Aquatic invertebrates: n = 78, μ = 485.3 nm; Aquatic vertebrates: n = 273 μ = 473.0 nm; Terrestrial invertebrates: n = 43, μ = 430.1 nm; Terrestrial vertebrates: n = 39, μ = 456.1 nm. B. Longest opsin: Aquatic invertebrates: n = 78, μ = 503.3 nm; Aquatic vertebrates: n = 273, μ = 507.6 nm; Terrestrial invertebrates: n = 43, μ = 530.3 nm; Terrestrial vertebrates: n = 39, μ = 539.1 nm. C. Opsin range: Aquatic invertebrates: n = 78, μ = 18.0 nm; Aquatic vertebrates: n = 273, μ = 34.6 nm; Terrestrial invertebrates: n = 43, μ = 100.2 nm; Terrestrial vertebrates: n = 39, μ = 83.0 nm. *: p < 0.05.

### (b) Terrestrial species saw shorter wavelengths of light than aquatic species

Terrestrial species were maximally sensitive to shorter wavelengths of light than aquatic species, but there was a significant interaction between habitat and lineage: aquatic vertebrate species were more sensitive to short wavelengths than aquatic invertebrate species, but terrestrial invertebrate species were more sensitive to short wavelengths that terrestrial vertebrate species. Additionally, invertebrates trended towards seeing short wavelengths of light (GLM, n = 433: habitat p = 0.045, t = −2.012, lineage p = 0.051, t = 1.960, interaction: p = 2.34*10^-3^; t = −3.061; λ_max_ shortest short-wavelength terrestrial species: 442±79.2 nm, aquatic species: 476±39.3 nm; invertebrate species: 466±70.2 nm, vertebrate species: 471±41.3 nm) (figure 1b).

### (c) Terrestrial species and invertebrates saw a larger range of colours than aquatic species and vertebrates

Terrestrial species saw a larger range of colours than aquatic species. In addition, there was a significant interaction between habitat and lineage: aquatic invertebrates saw a narrower range of colours than aquatic vertebrates, but terrestrial invertebrates saw a broader range of colours than terrestrial vertebrates (GLM, n = 443: habitat p = 2.51*10^-6^, t = 4.772, lineage p = 0.03, t = −2.184, interaction: p = 0.00261; t =2.232; λ_max_ range terrestrial species: 92±85.6 nm, aquatic species: 30.9±51.6 nm; invertebrate species: 47.2±80.4 nm, vertebrate species: 40.7±56.7 nm) (figure 1c).

### (d) Accounting for phylogeny removes the effect of habitat and lineage on visual pigment sensitivity

When we ran our analyses with only the subset of species included in our phylogenetic tree, we found that we lost the effect of lineage on the range of visual pigment sensitivity but did not lose the effect of lineage on the longest or shortest wavelengths of maximum sensitivity (see the electronic supplementary materials: tables S2 and S3). With this in mind, we controlled for phylogeny. Controlling for phylogeny removed the effect of habitat and lineage on longest λ_max_, shortest λ_max_ and range of λ_max_ (table 1).

**Table 1.** No effect of habitat or lineage on visual sensitivity following phylogenetic control.

| Variable | *Longest $\lambda_{\max}$* |  |  | *Shortest $\lambda_{\max}$* |  |  | *$\lambda_{\max}$ Range* |  |  |
| --- | --- | --- | --- | --- | --- | --- | --- | --- | --- |
|  | *p* | *t* | *SE* | *p* | *t* | *SE* | *p* | *t* | *SE* |
| *Habitat* | 0.257 | -1.13 | 11.4 | 0.439 | 0.774 | 17.0 | 0.226 | -1.211 | 21.7 |
| *Lineage* | 0.953 | 0.058 | 12.5 | 0.940 | 0.075 | 18.5 | 0.978 | -0.028 | 23.7 |
| *Habitat \* Lineage* | 0.474 | -0.72 | 20.3 | 0.214 | -1.25 | 30.1 | 0.552 | 0.596 | 38.6 |

### (e) Forest-woodland and open habitat species have similar spectral sensitivities

There was no effect of tree canopy openness on λ_max_, shortest λ_max_, and range of λ_max_ (table 2; see the electronic supplementary materials: figure S2, table S4 and table S5).

**Table 2.**
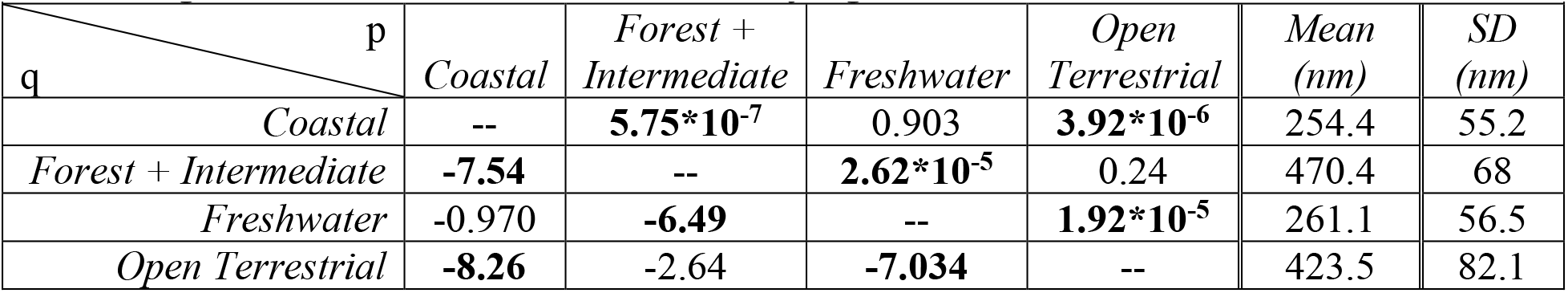
Effect of habitat greenness on shortest λ_max_. P-values are above diagonal; q-values are below diagonal. Numbers in bold are statistically significant.

### (f) Average and maximum depth, but not minimum depth, influenced sensitivity to blue but not red light

Species living at deeper average depth had longer shortest λ_max_ than shallow-living species (p = 0.046, t = 2.03; see the electronic supplementary material: figure S3a). Additionally, species living at a deeper maximum depth had shortest λ_max_ that were longer than shallow-living species shortest λ_max_ (p = 0.033, t = 2.18; electronic supplementary material: figure S4). Average depth and minimum depth did not affect species’ longest λ_max_ or range of λ_max_, and there was no effect of maximum depth on spectral sensitivity (electronic supplementary material: figures S3b-c and S5). Controlling for phylogeny removes the effect of average depth on shortest λ_max_ (phylolm: t = −4.688*10^-1^, p = 0.641).

### (g) Animals in coastal and freshwater habitats saw shorter wavelengths while animals in forest+intermediate or open-canopy habitats saw longer wavelengths

We found that coastal animals’ and freshwater animals’ shortest λ_max_ were shorter than both forest+intermediate animals’ shortest λ_max_ and open terrestrial animals’ shortest λ_max_ (omnibus test: p = 1.359*10^-12^, χ^2^ = 53.296, df = 3, pair-wise comparisons: table 2; figure 2). We also found that forest+intermediate animals’ longest λ_max_ and open terrestrial animals’ longest λ_max_ were longer than freshwater animals’ and coastal animals’ longest λ_max_ (omnibus test: p = 1.19*10^-14^, χ^2^ = 67.92, df = 3; pairwise comparisons: table 3 and figure 2).

**Table 3.** Effect of habitat greenness on longest λ_max_. P-values are above diagonal; q-values are below diagonal. Numbers in bold are statistically significant.

| p \ q | Coastal | Forest + Intermediate | Freshwater | Open Terrestrial | Mean (nm) | SD (nm) |
| --- | --- | --- | --- | --- | --- | --- |
| Coastal | -- | **$1.72 \times 10^{-7}$** | 0.77 | **$6.23 \times 10^{-10}$** | 297.3 | 72.4 |
| Forest + Intermediate | **-7.82** | -- | **$7.54 \times 10^{-6}$** | 0.22 | 547.1 | 17.7 |
| Freshwater | -2.40 | **-6.85** | -- | **$2.77 \times 10^{-5}$** | 314.3 | 73.8 |
| Open Terrestrial | **-9.14** | -2.71 | **-7.73** | -- | 534.1 | 49.1 |

**Figure 2.**
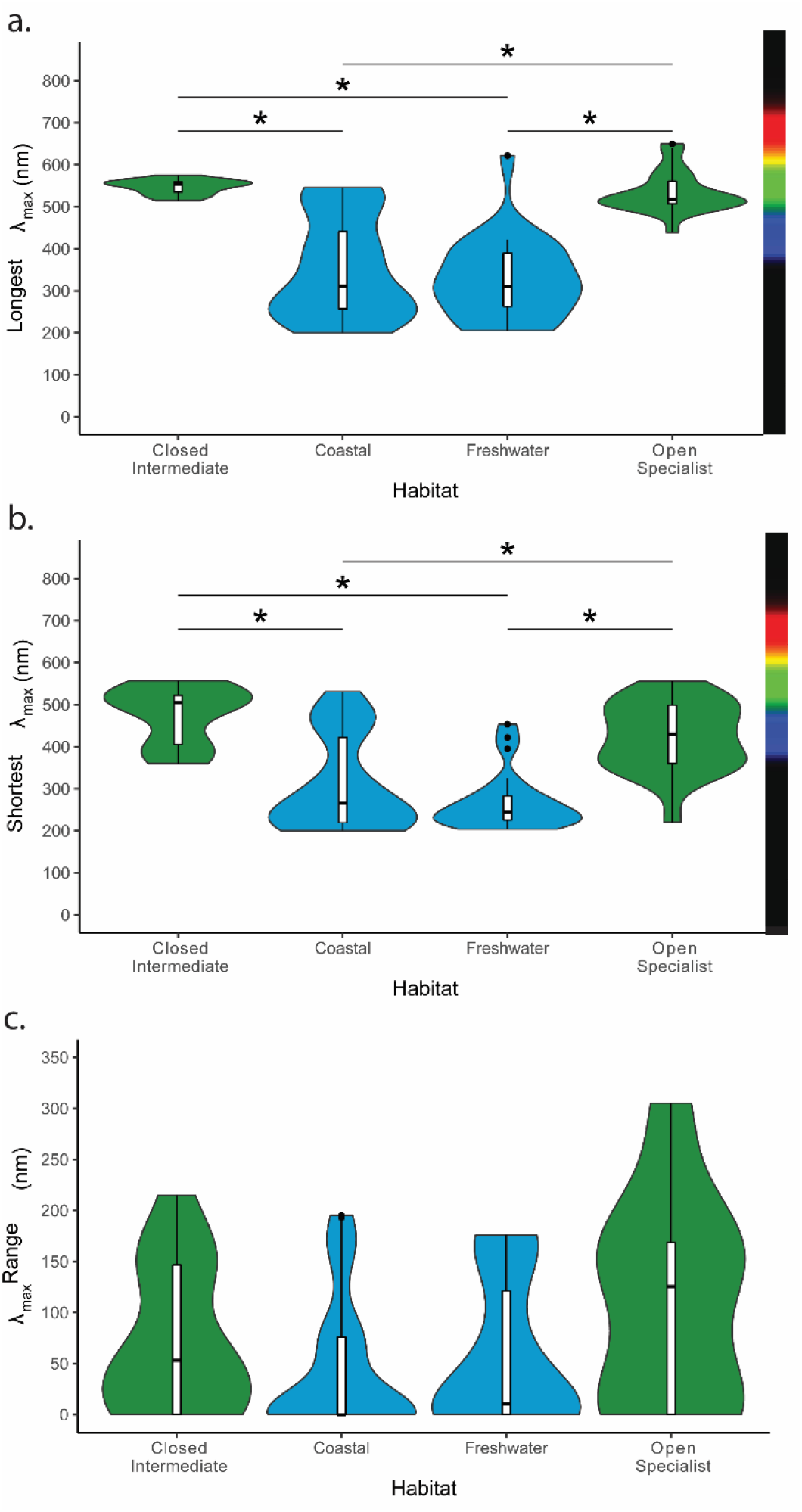
Effect of habitat greenness on visual sensitivities. A. Longest opsin: forest + intermediate: n = 14, μ = 547.1 nm, sd = 17.71 nm; coastal: n = 40, μ = 297.3 nm, sd = 72.42 nm; freshwater: n = 23, μ = 314.3 nm, sd = 73.77 nm; open terrestrial: n = 22, μ = 534.1 nm, sd = 49.09 nm; B. Shortest opsin: forest + intermediate: n = 14, μ = 470.4 nm, sd = 60.00 nm; coastal: n = 40, μ = 254.4 nm, sd = 55.16 nm; freshwater: n = 23, μ = 261.1 nm, sd = 56.52 nm; open terrestrial: n = 22, μ = 423.4 nm, sd = 82.06 nm C. Opsin range: forest + intermediate: n = 14, μ = 76.63 nm, sd = 72.61 nm; coastal: n = 40, μ = 742.88 nm, sd = 70.15 nm; freshwater: n = 23, μ = 53.26 nm, sd = 69.30 nm; open: n = 22, μ = 110.6 nm, sd = 98.82 nm. *: p < 0.05.

### (h) Open terrestrial animals had a broader visual range than coastal animals

We found that open terrestrial animals had a larger range (longest λ_max_ – shortest λ_max_) than coastal animals, but all other habitat groups were statistically similar (omnibus test: p = 0.03366, χ^2^ = 8.6931, df = 3; pairwise comparisons: table 4).

**Table 4.** Effect of habitat greenness on λ_max_ range. P-values are above diagonal; q-values are below diagonal. Numbers in bold are statistically significant.

| p \ q | Coastal | Forest + Intermediate | Freshwater | Open Terrestrial | Mean (nm) | SD (nm) |
| --- | --- | --- | --- | --- | --- | --- |
| Coastal | -- | 0.239 | 0.411 | **0.0497** | 42.88 | 70.2 |
| Forest + Intermediate | -2.65 | -- | 0.827 | 0.756 | 76.63 | 72.6 |
| Freshwater | -2.18 | -1.21 | -- | 0.561 | 53.26 | 69.3 |
| Open Terrestrial | **-3.64** | -1.40 | -1.84 | -- | 110.6 | 98.9 |

### (i) Accounting for lineage removed the effects of water and dissolved particles on visual pigment sensitivity

Controlling for phylogeny removed the effect of habitat greenness on shortest λ_max_, longest λ_max_, and λ_max_ range (table 5), even when we accounted for the species absent from our phylogenetic tree (see the electronic supplementary material: tables S6 – S8).

**Table 5.**
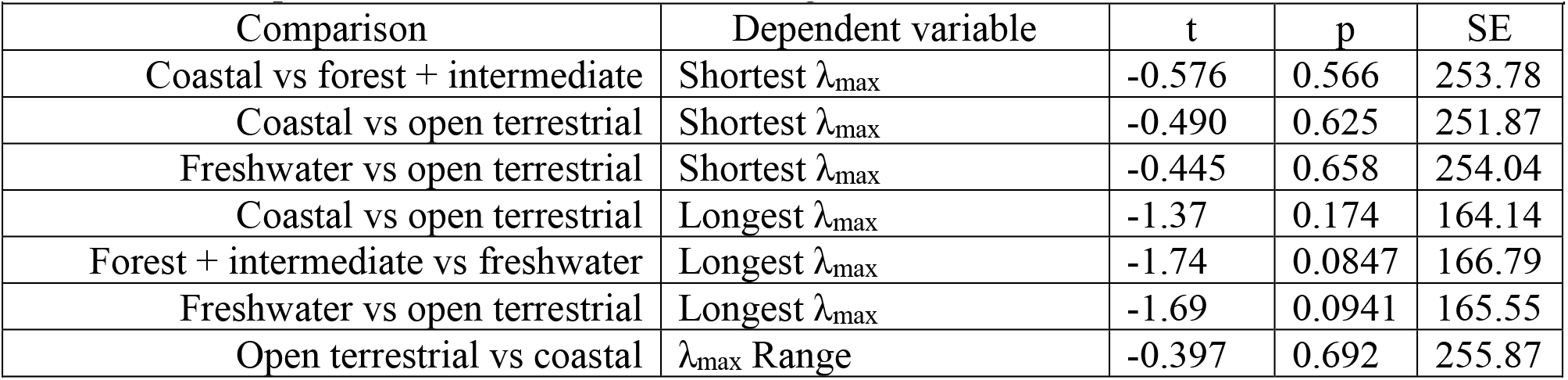
Loss of effect of habitat greenness on visual sensitivity following phylogenetic correction. Comparisons shown are those found significant in tables 2 – 4.

| Comparison | Dependent variable | t | p | SE |
| --- | --- | --- | --- | --- |
| Coastal vs forest + intermediate | Shortest $\lambda_{\max}$ | -0.576 | 0.566 | 253.78 |
| Coastal vs open terrestrial | Shortest $\lambda_{\max}$ | -0.490 | 0.625 | 251.87 |
| Freshwater vs open terrestrial | Shortest $\lambda_{\max}$ | -0.445 | 0.658 | 254.04 |
| Coastal vs open terrestrial | Longest $\lambda_{\max}$ | -1.37 | 0.174 | 164.14 |
| Forest + intermediate vs freshwater | Longest $\lambda_{\max}$ | -1.74 | 0.0847 | 166.79 |
| Freshwater vs open terrestrial | Longest $\lambda_{\max}$ | -1.69 | 0.0941 | 165.55 |
| Open terrestrial vs coastal | $\lambda_{\max}$ Range | -0.397 | 0.692 | 255.87 |

## (4) Discussion

### (a) The transition from aquatic to terrestrial habitats has influenced animal vision

We found that terrestrial species see longer long-wavelength light and a larger range of colours overall compared to aquatic species. Few other studies have broadly investigated the effect of animals’ evolutionary transitions between aquatic and terrestrial habitats on colour vision. However, transitions from aquatic to terrestrial life stages that lead to the development of different visual abilities can indicate whether differences between terrestrial and aquatic lifestyles themselves necessitate different strategies for perceiving the world [49]. Such studies have been conducted within single species: for example, in several species of dragonflies, adults have short wavelength-shifted vision, express more visual pigments than larvae, and have dorsal eye regions specialized to detect shorter wavelengths of light refracted from the sky [49,50]. Similar types of visual shifts have been observed in southern leopard frogs [66]. Just as animal development favours the expression of environmentally matched opsins over an intra-generational timescale, our results suggest that evolutionary adaptation favours the use of environmentally matched opsins over an inter-generational timescale.

The results of our terrestrial vs. aquatic models are congruent with the visual tuning hypothesis, that animal visual systems undergo adaptation to best detect the light most often present in their environments [23]. Terrestrial animals are exposed to a dynamic range of colours that changes throughout the day, including both short and long-wavelength light, as well as ultraviolet light in large forest gaps and open environments [25,26]. By contrast, aquatic animals, which we found to be less sensitive to long-wavelength and ultraviolet light, live in environments that are exposed to relatively less long-wavelength and ultraviolet light [24]. Absent phylogenetic controls, our regressions suggest that animals are likely to be maximally sensitive to colours most often present in their environment, and insensitive to colours likely to be absent.

### (b) Canopy coverage does not influence visual tuning

We found that animals which live in densely forested environments do not differ in their visual sensitivities from animals that live in open, prairie-like habitats. Although the forest light environment directly beneath the canopy is dominated by middle wavelengths (i.e., greens and yellows) under most conditions [25,26], spatial and temporal variations in forest light’s spectral qualities may require forest animals to possess visual sensitivities similar to those of animals living in open habitats.

Additionally, animals may choose to use light microhabitats which are suitable to their current visual physiology. Endler and Théry observed that forest birds use areas in which they are most conspicuous to advertise to potential mates [67]. Some species also modify their habitats to improve the visibility of their visual displays. For example, male golden-collared manakins clean the arenas they use to court females; the background of a cleaned arena contrasts better with male manakins’ plumage than the background of the forest surrounding the arena [68]. Arena cleaning also seems to improve white-bearded manakins’ ability to detect predators [69]. In such cases, evolution may be driving site preferences which match vision rather than driving vision to match site preferences, a complete reversal of the mechanism being investigated in our study.

### (c) The ciliary/rhabdomeric opsin divergence may impact the colours that animals can see

We found that animals that use rhabdomeric opsins for vision see a broader range of wavelengths of light than animals that use ciliary opsins for vision. Many animals that use rhabdomeric photoreceptors for vision, especially arthropods, have opsins that are maximally sensitive to ultraviolet light [31,70–75]. By contrast, comparatively few animals that use ciliary photoreceptors for vision have opsins that are maximally sensitive to UV light, although several species of birds and fish are sensitive to ultraviolet light [40,76–78]. Additionally, many mammals that utilize high acuity colour vision and whose short wavelength sensitive photoreceptors are sensitive to UV light, have corneas that selectively filter UV, inhibiting their ability to see those wavelengths [79,80]. Both ciliary and rhabdomeric opsins are thought to have been present in the urbilaterian, the common ancestor of all modern animals save sponges, cnidarians, placozoans, and ctenophores [55]. The emergence in chordates of ciliary opsins for vision rather than photoentrainment represents a singular event, one that may have also heralded differences in visual perception associated with reduced sensitivity to short wavelengths of light.

### (d) Phylogeny outweighs the effect of habitat

We found that the effects of habitat upon the spectrum of light animals can see were reduced once we controlled for phylogenetic history. These findings differ from those of studies looking at individual animal clades. For example, a 2018 survey of ray-finned fish found that species living at depth have reduced chromacy even after controlling for phylogeny [47]. Similarly, a historic study of cottoid fish in Lake Baikal found that there was a correlation between λ_max_ and habitat depth [81]. While studies of marine mammals found that species that forage near the surface have visual pigments that resemble those of terrestrial mammals while those that foraged at depth had visual pigments with amino acid substitutions that shifted the λ_max_ towards shorter wavelengths [82]. We found that, when expanding to include multiple clades – both chordates and non-chordates – a similar pattern emerged: terrestrial species had broader sensitivity to light and more sensitivity to long wavelengths of light compared to aquatic animals. However, these effects are lost once we account for phylogeny. This loss of an effect might be because the historical divergence between the visual pigments used by vertebrates and invertebrates is an important limiting factor on the degree to which visual pigments can accommodate for light environment, something that would not be detected in analyses limited to vertebrates.

The effect of the c-opsin/ r-opsin divergence is lost once we account for phylogeny in our analyses, but since this transition happened once and maps onto the metazoan phylogenetic tree, this loss of effect might be expected. Outside of this transition, opsin evolutionary history such as mutation biases may account for the effect of phylogeny on visual ability in our analysis. Retinal is covalently bonded to opsin *via* a Schiff base and the charge of the amino acid residues near the Schiff base influence the ability of retinal to change conformation and λ_max_ of the associated opsin [83,84], which has been experimentally confirmed using directed mutagenesis [37,85,86]. Future research should consider whether there are inherent differences in the electronic charge of the binding pocket between ciliary and rhabdomeric type opsins. Additionally, studies examining whether non-opsin means of visual tuning, including the differential absorption of light by screening pigments, differ between animals which use ciliary and rhabdomeric opsins and which live in the same light environment may prove particularly illuminating.

### (e) Conclusions

Here we used visual sensitivity data from nearly 450 animal species and 3 phyla to conduct a systematic survey of the effects of habitat light on the colours animals can see. We found that terrestrial animals and aquatic animals possess different ranges of spectral sensitivity from each other, but that evolutionary processes such as the c-opsin/r-opsin divergence may have limited chordates’ ability to tune their opsins to short-wavelength light. Additionally, the eyes of animals living in terrestrial habitats are not specifically tuned to forest canopy cover. Future research should consider whether inherent differences between chordate and non-chordate opsin amino acid sequences, or downstream neural signalling, are responsible for the evolutionary limitations to visual tuning.

## Acknowledgments

We would like to thank Adam Siepielski for training in systematic analysis, Jeremy Beaulieu for advice on conducting phylogenetically weighted regressions, and Nagayasu Nakanishi for his valuable feedback on early versions of this project and manuscript.

## Data Accessibility

Data pertaining to this study are available in the supplementary materials or available at Dryad repository https://doi.org/10.5061/dryad.47d7wm3fc. Code pertaining to this study is available at GitHub repository: https://github.com/mjosmurphy/opsin-evolutionary-ecology.

