## Supplementary Information for "Evolutionary history limits species’ ability to match color sensitivity to available habitat light"

M.J. Murphy[1,*] & E.L. Westerman[1]

[1]Department of Biological Sciences, University of Arkansas, Fayetteville AR 72701, USA

**Contents**

**Table S1:** Excel spreadsheet of species data used, with references.
**Table S2:** Excluding species flagged as *incertae cedis* and/or without sequencing data does not influence relationship between longest $\lambda_{max}$, shortest $\lambda_{max}$ or $\lambda_{max}$ range and coarse terrestrial habitat or coarse lineage.
**Table S3:** Sample size of combinations of independent variables in coarse habitat and coarse lineage analysis, before and after excluding species listed as *incertae cedis* and/or without sequencing data.
**Table S4:** Excluding species flagged as *incertae cedis* and/or without sequencing data does not influence relationship between longest $\lambda_{max}$, shortest $\lambda_{max}$ or $\lambda_{max}$ range and canopy cover.
**Table S5:** Sample size of animals living in each habitat in terrestrial animal analysis, before and after excluding species listed as *incertae cedis* and/or without sequencing data.
**Table S6:** Excluding species that were *incertae cedis* and/or without sequencing data did not influence the relationship between habitat greenness and longest or shortest $\lambda_{max}$, but did influence the relationship between habitat greenness and $\lambda_{max}$ range.
**Table S7:** Steel-Dwass post-hoc tests, following Kruskal-Wallis, examining the effect of habitat greenness on mean visual pigment sensitivities, prior to phylogenetic control, and prior to excluding species that were *incertae cedis* and/or without sequencing data.
**Table S8:** Sample size of animals living in each habitat in habitat greenness analysis, before and after excluding species listed as *incertae cedis* and/or without sequencing data.
**Figure S1:** Cladogram of phyla used in phylogenetic control
**Figure S2:** Terrestrial habitat type does not affect opsins' wavelength of maximum sensitivity
**Figure S3:** Species' average depth in the water column influences their shortest, but not longest or range in $\lambda_{max}$.
**Figure S4:** Species' minimum depth in the water column does not influence their, longest or range in $\lambda_{max}$
**Figure S5:** Species' maximum depth in the water column influences their shortest, but not longest or range in $\lambda_{max}$.
**Supplemental References**

**Table S1.** Excel spreadsheet of species data used, with references

(Not included with preprint. All data collected for this manuscript was taken from the Supplemental References, below.

See Dryad repository https://datadryad.org/stash/dataset/doi:10.5061/dryad.47d7wm3fc for processed data, when available)

**Table S2.** Excluding species flagged as *incertae cedis* and/or without sequencing data does not influence relationship between longest $\lambda_{max}$, shortest $\lambda_{max}$ or $\lambda_{max}$ range and coarse terrestrial habitat or coarse lineage. Significant comparisons in boldface.

| | *Before Excluding Data* | | | *After Excluding Data* | | |
|---|---|---|---|---|---|---|
| | **SE** | **t** | **p** | **SE** | **t** | **p** |
| ***Longest*** | | | | | | |
| *Habitat* | **5.64** | **5.60** | **$3.83*10^{-8}$** | **5.72** | **5.65** | **$3.00*10^{-8}$** |
| *Lineage* | 4.23 | -1.02 | 0.309 | 4.33 | -0.377 | 0.706 |
| *Habitat * Lineage* | 8.42 | -0.532 | 0.595 | 8.53 | -0.827 | 0.409 |
| ***Shortest*** | | | | | | |
| *Habitat* | **8.84** | **-2.01** | **$4.48*10^{-2}$** | **8.39** | **-2.02** | **$4.37*10^{-2}$** |
| *Lineage* | 6.29 | 1.96 | 0.0507 | 6.35 | 1.65 | 0.0990 |
| *Habitat * Lineage* | **12.5** | **-3.06** | **$2.34*10^{-3}$** | **12.5** | **-3.33** | **$9.49*10^{-4}$** |
| ***Range*** | | | | | | |
| *Habitat* | **10.1** | **4.77** | **$2.51*10^{-6}$** | **10.2** | **4.83** | **$1.91*10^{-6}$** |
| *Lineage* | **7.61** | **-2.18** | **$2.95*10^{-2}$** | 7.72 | -1.57 | 0.117 |
| *Habitat * Lineage* | **15.2** | **2.23** | **$2.61*10^{-2}$** | **15.2** | **2.28** | **$2.35*10^{-2}$** |

**Table S3**. Sample size of combinations of independent variables in coarse habitat and coarse lineage analysis, before and after excluding species listed as *incertae cedis* and/or without sequencing data.

| | | $n_{total}$ | $n_{excluded}$ | $n_{used\_in\_phylolm}$ |
|---|---|---|---|---|
| *Aquatic* | *Invertebrate* | 78 | 3 | 75 |
| | *Vertebrate* | 273 | 26 | 247 |
| *Terrestrial* | *Invertebrate* | 43 | 1 | 42 |
| | *Vertebrate* | 39 | 1 | 38 |

**Table S4.** Excluding species flagged as *incertae cedis* and/or without sequencing data does not influence relationship between longest $\lambda_{max}$, shortest $\lambda_{max}$ or $\lambda_{max}$ range and canopy cover.

| *Dependent variable* | *Before Excluding Data* | | | *After Excluding Data* | | |
|---|---|---|---|---|---|---|
| | **$\chi^2$** | **df** | **p** | **$\chi^2$** | **df** | **p** |
| *Longest $\lambda_{max}$* | 4.9986 | 2 | 0.08214 | 4.7425 | 2 | 0.09336 |
| *Shortest $\lambda_{max}$* | 4.3746 | 2 | 0.1122 | 3.5176 | 2 | 0.1723 |
| *$\lambda_{max}$ Range* | 1.1827 | 2 | 0.5536 | 0.89993 | 2 | 0.6376 |

**Table S5.** Sample size of animals living in each habitat in terrestrial animal analysis, before and after excluding species listed as *incertae cedis* and/or without sequencing data.

| | $n_{total}$ | $n_{excluded}$ | $n_{used\_in\_phylolm}$ |
|---|---|---|---|
| *Forest + Intermediate* | 15 | 1 | 14 |
| *Generalist* | 32 | 0 | 32 |
| *Open Specialist* | 22 | 0 | 22 |

**Table S6.** Excluding species that were *incertae cedis* and/or without sequencing data did not influence the relationship between habitat greenness and longest or shortest $\lambda_{max}$, but did influence the relationship between habitat greenness and $\lambda_{max}$ range.

| *Dependent variable* | *Kruskal-Wallis, Before Excluding Data* | | | *Kruskal-Wallis, After Excluding Data* | | |
|---|---|---|---|---|---|---|
| | $\chi^2$ | **df** | **p** | $\chi^2$ | **df** | **p** |
| *Longest $\lambda_{max}$* | 52.736 | 3 | $2.087*10^{-11}$ | 68.819 | 3 | $7.64*10^{-15}$ |
| *Shortest $\lambda_{max}$* | 41.553 | 3 | $4.993*10^{-9}$ | 58.296 | 3 | $1.359*10^{-12}$ |
| *$\lambda_{max}$ Range* | 7.3515 | 3 | 0.0615 | **8.5126** | **3** | **0.03652** |

**Table S7.** Steel-Dwass post-hoc tests, following Kruskal-Wallis, examining the effect of habitat greenness on mean visual pigment sensitivities, prior to phylogenetic control, and prior to excluding species that were *incertae cedis* and/or without sequencing data.

| | *Coastal* | *Forest + Intermediate* | *Freshwater* | *Open Terrestrial* | *Mean (nm)* | *SD (nm)* |
|---|---|---|---|---|---|---|
| **Longest** | | | | | | |
| *Coastal* | -- | **$8*10^{-7}$** | 0.93 | **$8.1*10^{-6}$** | 353.68 | 118.23 |
| *Forest + Intermediate* | **-7.46** | -- | **$1.1*10^{-5}$** | 0.20 | 547.54 | 17.16 |
| *Freshwater* | -0.855 | **-6.74** | -- | **$4.8*10^{-7}$** | 327.15 | 95.62 |
| *Open Terrestrial* | **-6.83** | -2.80 | **-7.59** | -- | 534.09 | 49.09 |
| **Shortest** | | | | | | |
| *Coastal* | -- | **$3.9*10^{-5}$** | 0.77 | **$4.5*10^{-4}$** | 309.58 | 109.38 |
| *Forest + Intermediate* | **-6.37** | -- | **$7.3*10^{-6}$** | 0.16 | 476.02 | 68.99 |
| *Freshwater* | -1.37 | **-6.86** | -- | **$1.2*10^{-5}$** | 269.10 | 67.79 |
| *Open Terrestrial* | **-5.59** | -2.96 | **-6.72** | -- | 423.45 | 82.06 |

| **Table S8.** Sample size of animals living in each habitat in habitat greenness analysis, before and after excluding species listed as *incertae cedis* and/or without sequencing data. | | | |
|---|---|---|---|
| | $n_{total}$ | $n_{dropped}$ | $n_{used\_in\_phylolm}$ |
| *Coastal* | 53 | 13 | 40 |
| *Forest + Intermediate* | 15 | 1 | 14 |
| *Freshwater* | 24 | 1 | 23 |
| *Open Terrestrial* | 22 | 0 | 22 |

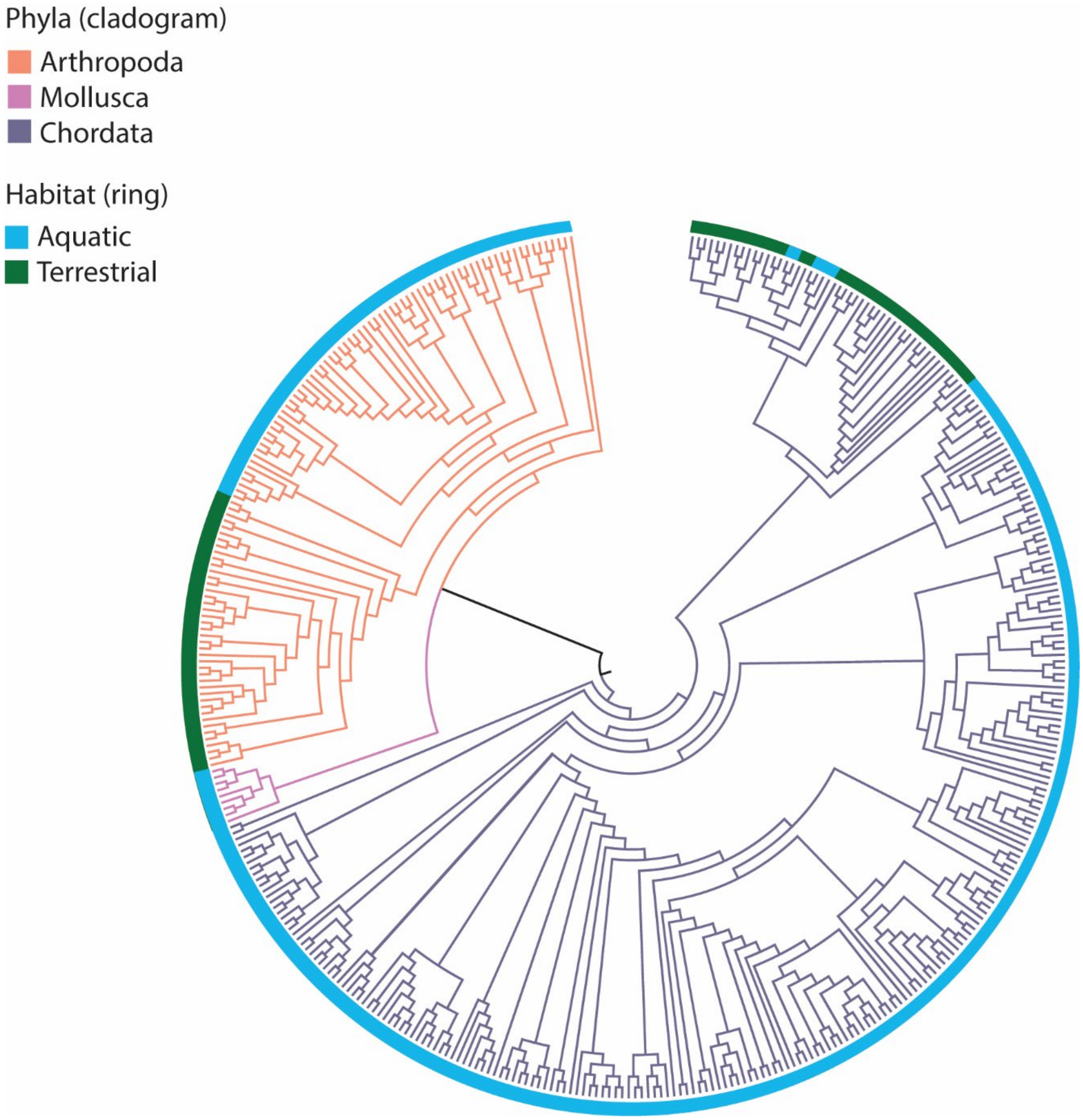


**Figure S1.** Cladogram of phyla used in phylogenetic control (n = 408 species).

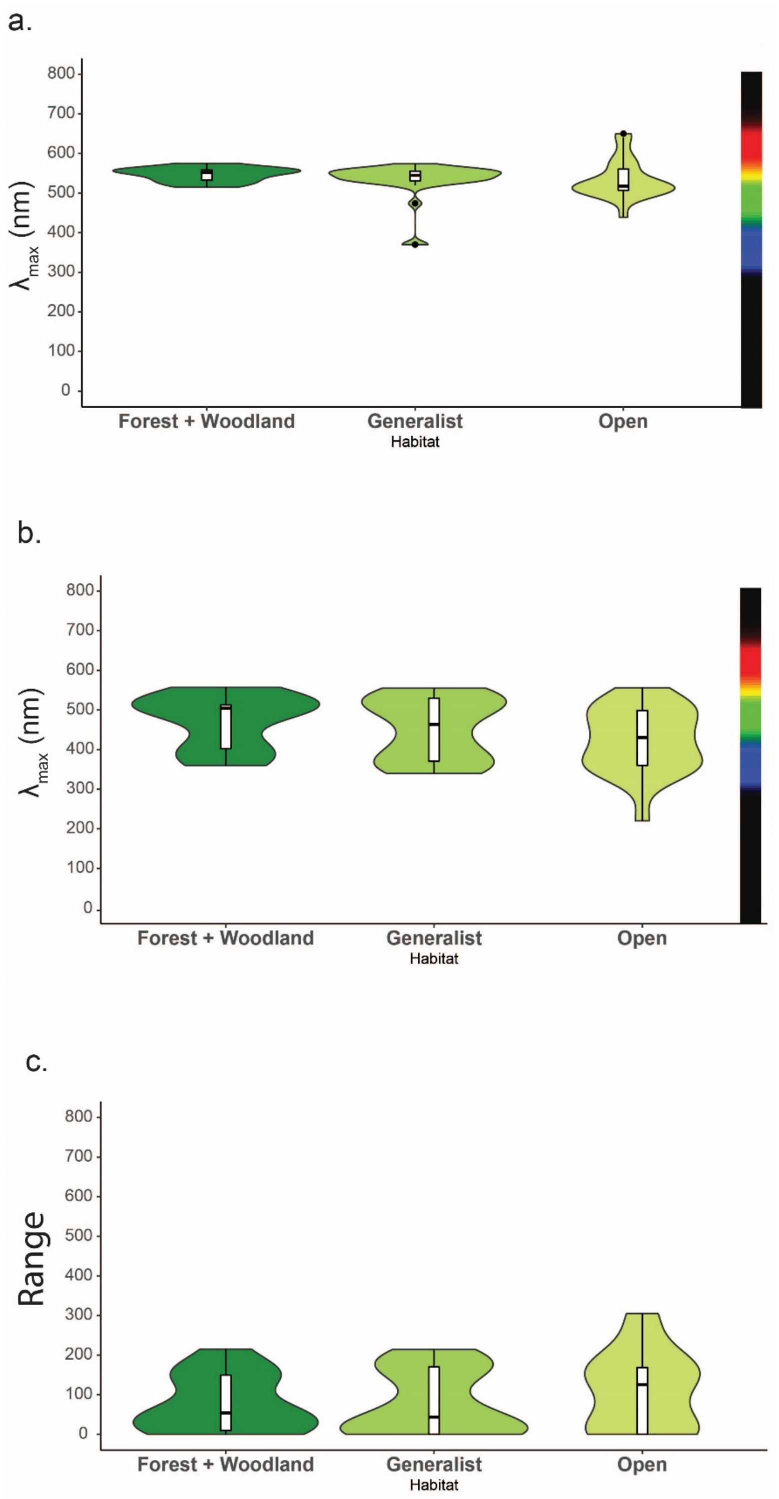


**Figure S2.** Terrestrial habitat type does not affect opsins' wavelength of maximum sensitivity; A. Longest opsin: forest + woodland: n = 14, μ = 546.9 nm; generalist n = 32, μ = 532.5 nm, open: n = 22, μ = 534.0 nm; B. Shortest opsin: Forest + woodland: n = 14, μ = 470.4 nm; generalist: n = 32, μ = 451.5 nm; open: n = 22, μ = 423.4 nm; C. Opsin Range: Forest + woodland: n = 14, μ = 76.5 nm; generalist: n = 32, μ = 80.9 nm; open: n = 22, μ = 111 nm. *: $p < 0.05$.

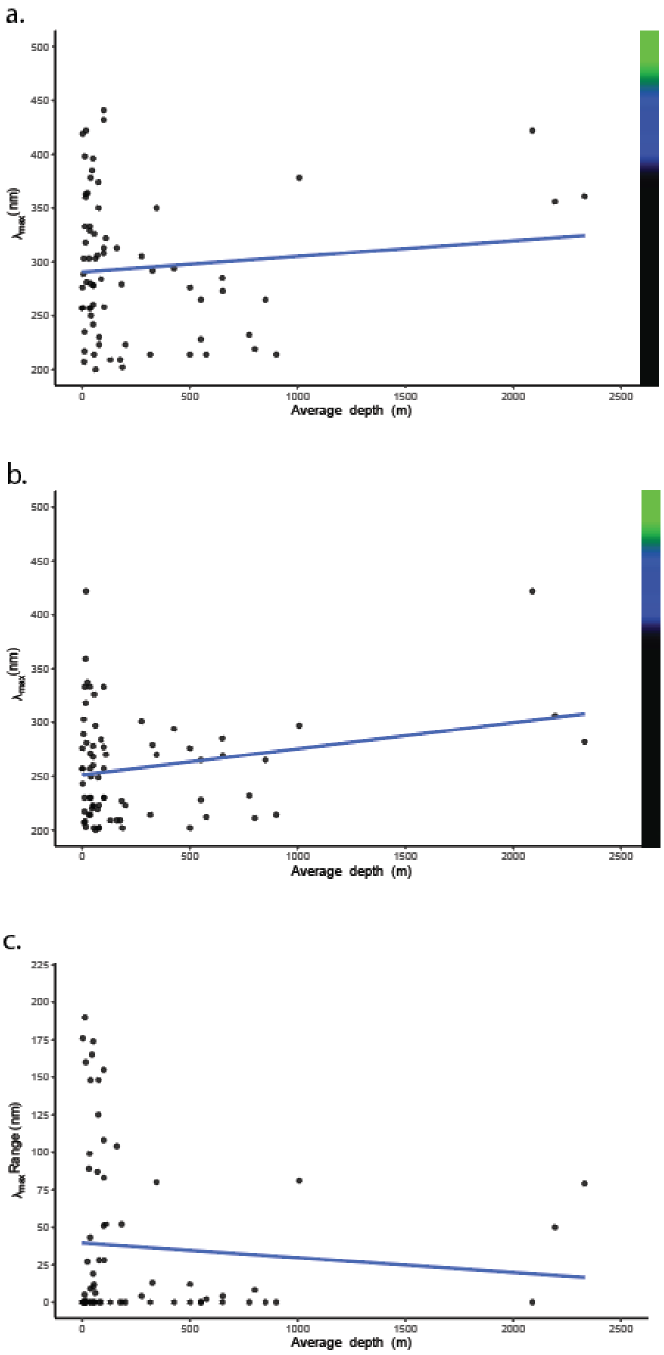


**Figure S3.** Species' average depth in the water column influences their shortest, but not longest or range in $\lambda_{max}$. A. Longest opsin: t = 0.90, p = 0.371, df = 72, SE = 0.016, AIC = 820.05; B. Shortest opsin: t = 2.02, p = 0.046, df = 72, SE = 0.012, AIC = 777.1; C. Range of opsins: t = -0.707, p = 0.482, df = 72, SE = 0.014, AIC = 798.9

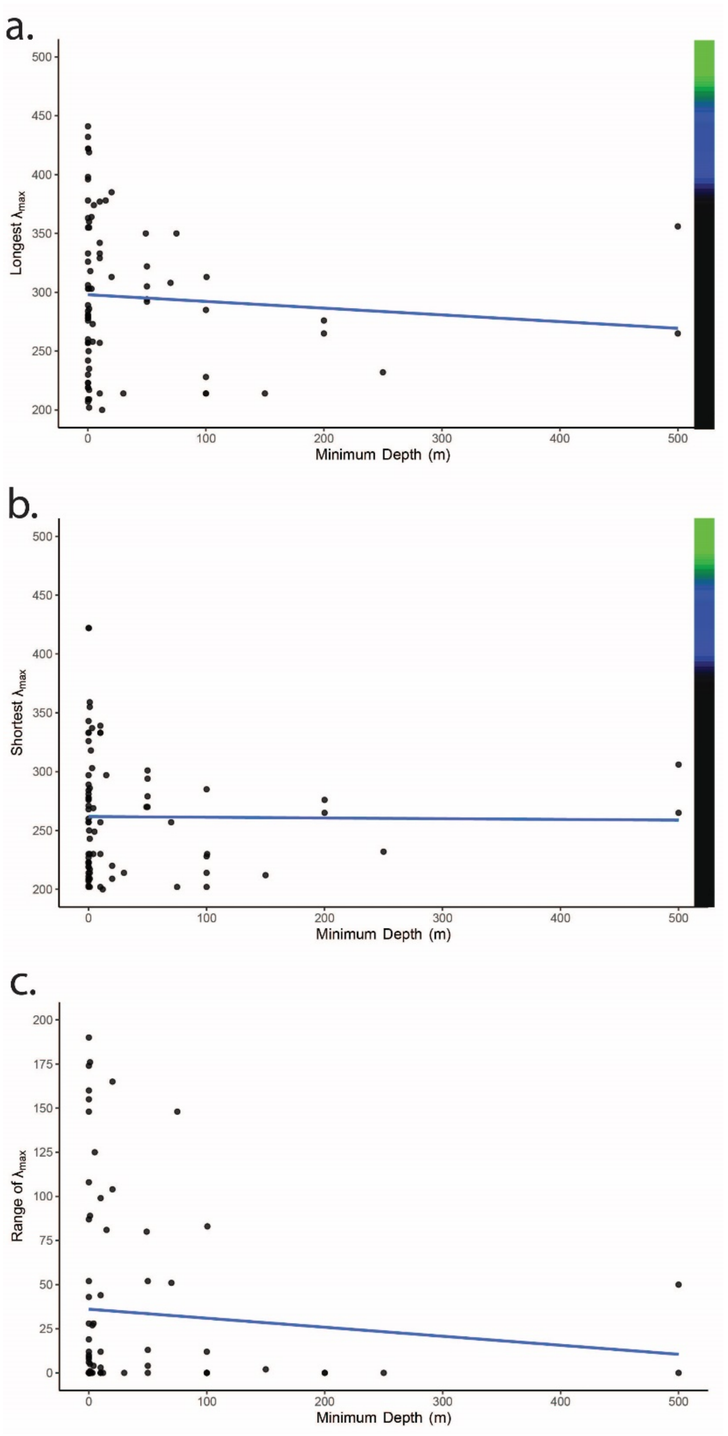


**Figure S4.** Species' minimum depth in the water column does not influence their, longest or range in $\lambda_{max}$. A. Longest opsin: $t = 0.059$, $p = 0.95$, $df = 78$, $SE = 0.0642$, $AIC = 887.1$; B. Shortest opsin: $t = 0.17$, $p = 0.87$, $df = 78$, $SE = 0.0511$, $AIC = 851.1$; C. Range of opsins: $t = -0.088$, $p = 0.93$, $df = 78$, $SE = 0.054$, $AIC = 860.3$

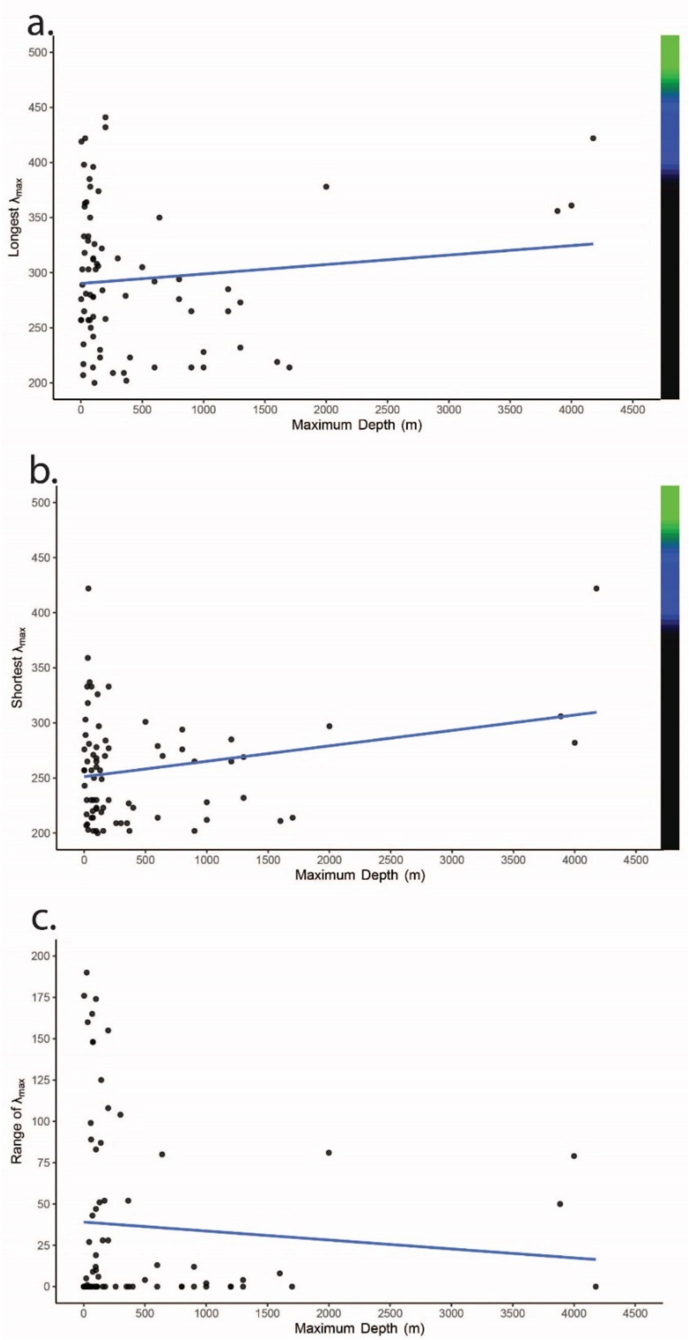


**Figure S5.** Species' maximum depth in the water column influences their shortest, but not longest or range in $\lambda_{max}$. A. Longest opsin: t = 0.99, p = 0.32, df = 73, SE = 0.0087, AIC = 840.5; B. Shortest opsin: t = 2.18, p = 0.033, df = 73, SE = 0.0064, AIC = 795.7; C. Range of opsins: t = -0.72, p = 0.47, df = 73, SE = 0.0075, AIC = 819.1

80. Van Hazel I, Sabouhanian A, Day L, Endler JA, Chang BS. 2013 Functional characterization of spectral tuning mechanisms in the great bowerbird short-wavelength sensitive

visual pigment (SWS1), and the origins of UV/violet vision in passerines and parrots. BMC Evol. Biol. 13. (doi:10.1186/1471-2148-13-250)

106. Beatty DD. 1969 Visual pigments of three species of cartilaginous fishes. Nature 222, 285.

121. Muntz WRA, Church E, Dartnall HJA. 1973 Visual pigment of the freshwater stingray, Paratrygon motoro. Nature 246, 517.

145. Chung W-S, Marshall NJ. 2016 Comparative visual ecology of cephalopods from different habitats. Proc. R. Soc. B 283, 20161346.
